## Supplementary File 1 for "The biogenesis of CLEL peptides involves several processing events in consecutive compartments of the secretory pathway"

**Supplementary File 1: Primers for cloning; restriction sites are underlined**

| **Name** | **5´- 3´ orientation** | **construct** | **Restriction sites** |
| --- | --- | --- | --- |
| EPI10 -1 F EcoRI | CCGAATTCATGTCTAGTAGCTTCCTTAG | EPI10-FLAG | EcoRI |
| EPI10 -711 R KDEL BamHI | CCGGATCCTTACAGCTCGTCCTTTTTGTCGTCATCATCCTTGTAGTC | EPI10-FLAG | BamHI |
| CLEL9 Prom F1 | AATTGCGGCCGCGAATCTATCACATCTCCCTCTTGACT | CLEL9:EPI1a/10 | NotI |
| CLEL9 Prom R1 | AATTGAATTCGACTTCTCTTATTCTTCTTCTTGTTCT | CLEL9:EPI1a/10 | EcoRI |
| CLEL6 Prom F1 | AATTGCGGCCGCATAAAATCTTGTTTCATTTG | CLEL6:EPI1a/10 | NotI |
| CLEL6 Prom R1 | AATTGAATTCGACTTCTCTTATTCTTCTTCTTGTTCT | CLEL6:EPI1a/10 | EcoRI |
| EPI1a F1 | AATTGAATTCATGAGTTCTAGTTTCTTGTCTAG | Flag-EPI1a | EcoRI |
| EPI1a F2 | GACTATAAGGATGATGATGATAAACAATCACCTCAAGTTATATCACC | Flag-EPI1a | ---- |
| EPI1a R1 | TTTATCATCATCATCCTTATAGTCACTTGAAGAAACGTGACAAAAA | Flag-EPI1a | ---- |
| EPI1a R2 | AATTCTCGAGTTACTTGCTAGGTGGCTGTTC | Flag-EPI1a | XhoI |
| EPI10 F1 | AATTGAATTCATGTCTAGTAGCTTCCTTAGTT | EPI10-Flag | EcoRI |
| EPI10 F2 | AATTCTCGAGTCATTTGTCGTCATCATCCTT | EPI10-Flag | XhoI |
| SPCLEL6 F3 to GFP | GTCTTCCAATGTTGGATGTGCCTCTGCCGTTAGCAAAGGTGAAGAAC | SP-sfGFP |  |
| SPCLEL6 F2 to GFP | GTGTTCTGTTTCATTCTTCTGCTTTTGTCTTCCAATGTTGGATG | SP-sfGFP |  |
| SPCLEL6 F1 to GFP | ATGTCTTGCTCTTTGAGGAGTGGACTCGTGATAGTGTTCTGTTTCATTCT | SP-sfGFP |  |
| sfGFP–Stopp BamHIR | GGATCCGCTGCCTTTATACAG |  |  |
| XylT35 F1 | ATGAGTAAACGGAATCCGAAGATTCTGAAGATTTTTCTGTATATGTTACT | XylT35-sfGFP |  |
| XylT35 F2 | TCTGTATATGTTACTTCTCAACTCTCTCTTTCTCATCATCTACTTCGTTT | XylT35-sfGFP |  |
| XylT35 to sfGFP F | TCATCTACTTCGTTTTTCACTCATCGTCGTTTTCAGTTAGCAAAGGTGAA | XylT35-sfGFP |  |
| XylT35 F1 EcoRI | CCGAATTCATGAGTAAACGGAATCCGAA | XylT35-sfGFP | EcoRI |
| CLEL6 -1 EcoRI | AATTGAATTCATGTCTTGCTCTTTGAGGAG | sfGFP | EcoRI |
| CLEL6 -401 BamH | AATTGGATCCAGACTTCTCGTTGTGGATCG | CLEL6 sec | BamHI |
| CLEL6 –88 BamHIF | GGATCCGCTCGTCGTTTAAGGTCAC | CLEL6 sec | BamHI |
| CLEL6 -88 R-A BamHIF | GGATCCGCTGCCGCCGCCGCCTCACATAAGC | CLEL6 R-A | BamHI |
| CLEL9 -79 BamHI F | GGATCCCGAAGCTTACGAGAAGTAG | CLEL9 sec | BamHI |
| CLEL9 -240 BamHIR | GGATCCTTAGCGATTGTGTATCGGTC | CLEL9 sec | BamHI |
| CLEL6 F 202 D-A | AGTAGTCATGGCTTATCCTCAGCCT | CLEL6 D71A | ---- |
| CLEL6 R 202 D-A | AGGCTGAGGATAAGCCATGACTACT | CLEL6 D71A | ---- |
| CLEL9 F 198 D-A | TTGATGGATATGGCTTATAATTCTGCG | CLEL9 D66A | ---- |
| CLEL9 R 198 D-A | CGCAGAATTATAAGCCATATCCATCAA | CLEL9 D66A | ---- |
| CLEL6 RRLR F1 | TGCCGCTGCCGCCTTAAGGTCACATAA | CLEL6 R-A | ---- |
| CLEL6 RRLR R1 | TTATGTGACCTTAAGGCGGCAGCGGCA | CLEL6 R-A | ---- |
| CLEL6 RRRAL F1 | TGGAGAAGCCGCCGCCGCGTTGGGTGGA | CLEL6 R-A | ---- |
| CLEL6 RRRAL R1 | TCCACCCAACGCGGCGGCGGCTTCTCCA | CLEL6 R-A | ---- |
| CLEL6 RRLR F2 | TGCCGCTGCCGCCGCCGCCTCACATAA | CLEL6 R-A | ---- |
| CLEL6 RRLR R2 | TTATGTGAGGCGGCGGCGGCAGCGGCA | CLEL6 R-A | ---- |
| CLEL6 RRRAL F2 | TGGAGAAGCCGCCGCCGCCGCCGGTGGA | CLEL6 R-A | ---- |
| CLEL6 RRRAL R2 | TCCACCGGCGGCGGCGGCGGCTTCTCCA | CLEL6 R-A | ---- |
| CLEL6-261 KDELR | GGATCCTTACAGCTCGTCCTTAGACTTCTCGTTGTGG | CLEL6 KDEL | BamHI |
| CLEL9 -240 KDELR | GGATCCTTACAGCTCGTCCTTGCGATTGTGTATCGGTC | CLEL9 KDEL | BamHI |
| CLEL9 154 F BamHI | GGATCCCACGAAGGTGGAGGAAGTGA | Δ-CLEL9 | BamHI |
| CLEL6 169 F BamHI | GGATCCGCGTTGGGTGGAGTCGAGAC | Δ-CLEL6 | BamHI |
| SBT3.8 F | GAATTCATGAAGAGTTGCAGAACCTT | SBT3.8 sfGFP | EcoRI |
| SBT3.8 F2 | ATGAAGAGTTGCAGAACCTT | SBT3.8 HIS | EcoRI |
| SBT3.6 HIS R | TCAGTGGTGGTGGTGGTGGTGGTTTTCATCGTAGTAGTTC | SBT3.8 HIS | EcoRI |
| SBT3.8 - Stopp | GGATCCGTTTTCATCGTAGTAGTTCTG | SBT3.8 sfGFP | BamHI |
| CLEL6 –SP NcoI F | AATTCCATGGGCGCTCGTCGTTTAAGGTCACA | Rec CLEL6 | NcoI |
| CLEL6 HIS NcoI R | AATCCATGGTCAGTGGTGATGATGGTGATGAGACTTCTCGTTGTGGATCG | Rec CLEL6 | NcoI |
