## Supplementary File 2 for "The biogenesis of CLEL peptides involves several processing events in consecutive compartments of the secretory pathway"

**Supplementary File 2: GreenGate modules used for this study including (A) sources of templates, primers used for cloning and (B) a general green gate module list**

**A**

| **Name** |  | **source of plasmid** | **PCR template** | **FW Oligo** | **RW Oligo** | **comment** |
| --- | --- | --- | --- | --- | --- | --- |
| pGGA-VHP1 | 3142 bp of VHP1 (At1g15690) promoter | this work | *Arabidopsis thaliana* genomic DNA | aacaGGTCTCAACCTgaagaattacgttaagctttaaag | aacaGGTCTCACGTCtcaccgacacaagtcc | amplified in two fragments to remove one internal Eco31I site |
|  |  |  |  | aacaGGTCTCAACCTcttaccgtgtaaaatcattcg | aacaGGTCTCATGTTcttctctcctccgtataagag |  |
| pGGB003 | B-Dummy | Lampropoulos et al., 2013 |  |  |  |  |
| pGGC-ManI | α-Mannosidase I (At1g51590) full length CDS | this work | *Arabidopsis thaliana* cDNA | aacaGGTCTCAGGCTcaacaatggcgagaagtagatcgat | aacaGGTCTCTCTGAaacgttaatctgatgaccaaact |  |
| pGGC-ST | N-terminus of rat sialyltransferase | this work | *P16_Pro_-ST-pHusion* (Luo et al. 2015) | aacaGGTCTCAGGCTcaacaatgattcataccaacttgaa | aacaGGTCTCTCTGAcatggccactttctcctggc |  |
| pGGD010 | mCherry | Schuster et al. 2014 |  |  |  |  |
| pGGE001 | rbcS terminator | Lampropoulos et al., 2013 |  |  |  |  |
| pGGF012 | *MAS_Pro_:SulfR_t35S* | Lampropoulos et al., 2013 |  |  |  |  |
| pGGZ003 | Destination vector | Lampropoulos et al., 2013 |  |  |  |  |

**B**

| **GreenGate module list** | |
| --- | --- |
| ***VHP1_Pro_:ManI-mCherry*** | |
| entry module | insert |
| pGGA-VHP1 | VHP1 promoter |
| pGGB003 | B-Dummy |
| pGGC-ManI | α-Mannosidase I |
| pGGD010 | mCherry |
| pGGE001 | rbcS terminator |
| PGGF012 | Sulfadiazine resistance |
| pGGZ003 | Destination vector |
| ***VHP1_Pro_:ST-mCherry*** | |
| entry module | insert |
| pGGA-VHP1 | VHP1 promoter |
| pGGB003 | B-Dummy |
| pGGC-ManI | N-terminus of rat sialyltransferase |
| pGGD010 | mCherry |
| pGGE001 | rbcS terminator |
| PGGF012 | Sulfadiazine resistance |
| pGGZ003 | Destination vector |
