## Supplementary File 3 for "The biogenesis of CLEL peptides involves several processing events in consecutive compartments of the secretory pathway"

**Supplementary File 3: Primers used for qPCR analysis**

| SBT1.1 F | CGTGCGATGAAGGCACAAACTTCT |
| --- | --- |
| SBT1.1 R | CCCAACAAGCTTTGTAAGCAGCGA |
| SBT1.2 F | TCAAACGTGGCAAGACTACGGAGA |
| SBT1.2 R | GTCAACTGCCCTTGTGCAAAGCTA |
| SBT1.3 F | AGCAGCTCCTTCATCGCCTTATGA |
| SBT1.3 R | ACAAAGCTGATATCGCCGGGTAGT |
| SBT1.4 F | TGAAACGAGCAAGCTAAGAACCGC |
| SBT1.4 R | GCCGAGCACAACGCTCTTAAATGT |
| SBT1.5 F | AGGCGATTCGGATTCGGTTTACGA |
| SBT1.5 R | AAGAGTGACCACCAAGGGACTTGT |
| SBT1.6 F | AGAGGAGTGACGGTGACAGTGAAA |
| SBT1.6 R | ACGTATCCATTTGGGTCACCACGA |
| SBT1.7 F | CGTACTCGGTCGCTGATTTGAACT |
| SBT1.7 R | ACGTGTTTCCCATCCGACCATTCA |
| SBT1.8 F | TTGGCTGGTTGGTCTGACGCTATT |
| SBT1.8 R | TTAGATAGGCTGTTGTCTGCAGCATCATG |
| SBT1.9 F | TCACGTCTGATCAAAGCAGTCCCA |
| SBT1.9 R | CCACCACAGAACAGCTCACTTCAA |
| SBT2.1 F | TGAGCATATAATGGCGCAACGCAC |
| SBT2.1 R | AGAGGCTGCAAGGGAAGAGTTGTA |
| SBT2.2 F | TAAAGGCGAGGCGATAATGGCTCA |
| SBT2.2 R | TCCGCTGATTGTTGCATTGTTCCG |
| SBT2.3 F | CCTTCAACAATCTCATCCGCGCTT |
| SBT2.3 R | TATCCGAGCCGTTGATCCCACAAA |
| SBT2.4 F | ACTTAAGCCAGATATTCTTGCACCTGGTC |
| SBT2.4 R | AACATGGCCCGCACCATGATCAAA |
| SBT2.5 F | TGGTCATGTCAATCCAAGTGCTGC |
| SBT2.5 R | TAGACGGCTGCATCCTTGCTGTTA |
| SBT2.6 F | AGGAAGACCTCTCCAAGCACAACA |
| SBT2.6 R | ATGAGAGATGGCTATGGATGGCGT |
| SBT3.1 F | GGAACATCAATGGCAACTCCGGTT |
| SBT3.1 R | TACAAGACCGGGATCTGTGGCTTT |
| SBT3.2 F | AAGAACGTTGGTTGGACTTCCCGT |
| SBT3.2 R | AATGAGGGTGCACAGACTTGAGGA |
| SBT3.3 F | ACATAGCAGCACCAGGAGTAAGGA |
| SBT3.3 R | ATATGAGACCGGGTTCTGCAGCTT |
| SBT3.4 F | TGCCGGTGGATTTGTTATGCGTTC |
| SBT3.4 R | TGAGACCTGGTTCTGCAGCTTTCT |
| SBT3.5 F | GTTGCGGGAGTTGTTGCACTTCTT |
| SBT3.5 R | TATGAGACCGGGATCTGCAGCTTT |
| SBT3.6 F | ATTGGCAGCTACAACCAACACCAC |
| SBT3.6 R | TCTGGATTTACAAGGCCTCCACCA |
| SBT3.7 F | GTTGACTACGAGCTAGGGACCTAT |
| SBT3.7 R | TGAGGATCGCTGCTGAAATTGGA |
| SBT3.8 F | AAGCTGGTGGTCTTGGCGTAATTG |
| SBT3.8 R | AATATGCTCACTCCTGGTGCTGCT |
| SBT3.9 F | CTCTGCTAATCCCAACAGTACAATGG |
| SBT3.9 R | CGTCTTTGAAGCCTGTATCTTTACAATG |
| SBT3.10 F | TGCCAATCCCAAGAGTGCAATG |
| SBT3.10 R | GTGTTCTTGAAGCTTGTATGTTTACAATGG |
| SBT3.11 F | TTTCAGGGAGAGGACCCAATTCCA |
| SBT3.11 R | TTAAGCGCAGCTGGAGACCAATCA |
| SBT3.12 F | ACATCCTATGCCACTCCCGTTGTT |
| SBT3.12 F | ATCTTTGGCTCTTTCCGCGTTCAC |
| SBT3.13 F | ACTGCAGCTACTACACTAACTGGTCA |
| SBT3.13 R | AGTCCAAATCCGTTCTGTTCTTCTGG |
| SBT3.14 R | CGGATTTGGACTTTATTCAGGGACATC |
| SBT3.14 F | TAGTCAAATGGATCTGCAAGCTTCT |
| SBT3.15 R | GCAATGTTGCTCCATGGCTCTTGA |
| SBT3.15 F | TGTGGCCGCACTAATTCTCACAGT |
| SBT3.16 F | CAGGTATACTCGCAGCCGTACCAA |
| SBT3.16 R | TACAAGGCCGCCTCCATAGTCAAA |
| SBT3.17 F | GCAGCACCGGGTGTTAATGTTCTT |
| SBT3.17 R | CGGGTGTGTTACTTTCTCTGGGTT |
| SBT3.18 F | AGCCAGATATTACAGCTCCTGGCA |
| SBT3.18 R | GGGTCCATGGCTTTCAAAGGGTTT |
| SBT4.1 F | ATCATAGCGGGTTGGCCTGAGAAT |
| SBT4.1 R | CTTGTCGCTTCCAACGTGTGATCT |
| SBT4.2 F | ATGTCGCAGGTGTAGCTGCTTATGTCAAG |
| SBT4.2 R | TGGATCAACTGCCACTGTAGGGTT |
| SBT4.3 F | CTTCCAGACAGCGCATTTGTGGTT |
| SBT4.3 R | TGAAGGAGAAGCTACCGGAGAAAATGCAG |
| SBT4.4 F | TTCACCTGACAGTTCACCGACTGA |
| SBT4.4 R | CCAGAACCAGAAGCATTCATTGGC |
| SBT4.5 F | AATGTCTTGTCCGCATGTTGCTGG |
| SBT4.5 R | TTTGCAGTGTAGTTCAAGCCGCAG |
| SBT4.6 F | TCTCCTTATGTCCCACCAAGCGAA |
| SBT4.6 R | AGCCGCAGAGAAAGGTGATATGGT |
| SBT4.7 F | TGTGGCTGCGTACATCAGGACTTT |
| SBT4.7 R | AGGCGATGTGGTCTGCTTTGTCTA |
| SBT4.8 F | TCCGGAACTTCTATGGCTTGTCCA |
| SBT4.8 R | TTCCACTGCAAATGACGGCATCAC |
| SBT4.9 F | GATGACTTTGACTCTGTCATCTCTTACGT |
| SBT4.9 R | AGCTGGTGAATTCAAAGGTGAAAACGCAG |
| SBT4.10 F | AGGAGTGGAAATCCTCGCTGCTTA |
| SBT4.10 R | AGTTCCAGAAGGATTCATCGGCCA |
| SBT4.11 F | GGCTGCAAATTCACTTAGGGCATC |
| SBT4.11 R | ATTGGGTCTACATGACCTGCTCCA |
| SBT4.12 F | GGTGGCTGCGTATGTCAAGACGTT |
| SBT4.12 R | TTGGAGCACTTGACGGTATCACCCGAGA |
| SBT4.13 F | TGCTGGTGAACCGTCTCAAGATGA |
| SBT4.13 R | AAGCAAACTCTGTCGATGCGATGC |
| SBT4.14 F | AATTCACCATCCTGTCTGGCACCT |
| SBT4.14 R | CTTCGCCGCACAAGAACTGAACAT |
| SBT4.15 F | TTGGTCTCCAGCCGCAATCAAATC |
| SBT4.15 R | GTGTTGTTGCTGTTATCGCCGGTT |
| SBT5.1 F | CAGAAATGCAGCACCAGAAGGCAA |
| SBT5.1 R | TAGTAACCTGCCCAGCACCAAAGT |
| SBT5.2 F | CATGGACTGGAAACGACTCAAGCA |
| SBT5.2 R | GGCTGTTGCACCAGTTTCTGTTGT |
| SBT5.3 F | AAGCCTGACATAACTGCTCCTGGT |
| SBT5.3 R | ACGAGACCGGGATTCACAGCTAAA |
| SBT5.4 F | CCTCATCAGTGCTGCAGATGCTA |
| SBT5.4 R | AAAGGAGCAGGCTTTGTGTTCAGC |
| SBT5.5 F | GCTGGTGCAATTGCCCTTCTCAAA |
| SBT5.5 R | TGCATCGTAGACTAAACCGGGAGA |
| SBT5.6 F | CAGGACTGTACATACTGGCTGCAT |
| SBT5.6 R | CCAAGTGCAAATGGGTTTGCTGGT |
| SBT6.1 F | ACTGCCATCCCGAAGAACTGATGT |
| SBT6.1 R | ACTCCACTAGCTACAACACCAGCA |
| SBT6.2 F | GGTGAGAAGGAAGGTGAAGGAGAA |
| SBT6.2 R | GGAAACGAACCTGCATCCATTGCT |
